# A global assessment reveals widespread co-benefits for forest protection

**DOI:** 10.64898/2026.10.22.760816

**Authors:** Charlotte H. Chang, Brian E. Robinson, Dávid Papp, István Molnár, James T. Erbaugh, Yuta J. Masuda

**Affiliations:** Dept. of Data Science, University of Oregon, USA; Dept. of Geography, McGill University, Montreal, Canada; International Center for Integrated Mountain Development, Kathmandu, Nepal; Budapest University of Technology and Economics, Hungary; Lexunit Zrt., Hungary; Global Science, The Nature Conservancy, USA; Dept. of Environmental Studies, Dartmouth College, USA; Allen Family Philanthropies, USA

## Abstract

Avoided forest conversion (“AFC”) is a high-impact conservation and climate mitigation strategy, yet its impacts on human well-being and ecosystems remain an open question. We present the first global and comprehensive assessment of AFC impacts on human material and non-material well-being, biodiversity, and ecosystem services in a set of 29,038 studies published between 1990-2022. We leveraged an ensemble of open- and closed-weight GPT models, training regimes, and architectures, and applied novel approaches to detect hallucination-induced reasoning errors. 85% of the studies report positive human and environmental co-benefits, with strong evidence for synergies rather than trade-offs. Nevertheless, there are evidence gaps for countries facing imminent forest conversion threats. Methodologically, this study advances an AI workflow for conducting complex, holistic evidence syntheses that traditionally require thousands of expert hours. The results reinforce AFC as an immediate climate and conservation solution capable of delivering substantial environmental and human well-being benefits.

## 1 Introduction

Protecting forests is critical for global biodiversity conservation and climate change mitigation goals (*1*–*6*), positioned at the core of ambitious area-based targets such as the Kunming-Montreal Global Biodiversity Framework (Target 3, or “30 *×* 30”, (*4, 7*)). As forests support the vast majority of global vertebrate and plant diversity (*5, 8*), avoided forest conversion (“AFC”) has emerged as one of the most high-impact and cost-effective pathways for near-term climate mitigation (*1, 9*–*11*). Successful AFC efforts have provided vital benefits for local communities, including health and livelihood improvements (*12*–*14*), and helped attract $84 billion USD in investment across all conservation sectors in 2023 (*15*). A primary assumption underpinning this substantial investment in AFC is that it yields net positive co-benefits. In practice, tensions, trade-offs, and implementation challenges have raised questions about the extent to which AFC delivers on these promises (*16, 17*).

Calls to comprehensively characterize the distribution of benefits versus trade-offs for AFC reveal that we do not know the true extent of evidence supporting or contesting AFC co-benefits (*16, 18*–*20*). This knowledge deficit is particularly important as forested ecosystems support more than one billion people (*21, 22*). Furthermore, AFC has been central to high profile debates focused on zero deforestation mandates, land or protected area management strategies, and benefit distribution models or payments for ecosystem services (*7, 23*–*26*). Research on these topics now span decades and systematic reviews have made some progress in providing a knowledge base of the benefits and trade-offs of AFC. For example, we now have a deeper understanding that co-benefits are often conditional on formal community governance (*27, 28*), but many claims remain speculative rather than empirically validated (*25, 29, 30*). Nonetheless, these reviews remain limited in scope due to the demands of scale and disciplinary complexity.

Any comprehensive survey of AFC co-impacts must contend with its inherently interdisciplinary scope and sheer volume (*31, 32*). This body of evidence encompasses over 120 disciplines (*33*), representing markedly different research designs, reporting structures, and analytical frameworks. The combined challenge of disciplinary breadth and volume has meant that impact estimates have historically relied on relatively small sample sizes derived from manual literature review or biophysical data proxies (e.g., *7, 29, 34, 35*). As a result, traditional meta-analyses of AFC pose stark trade-offs: they may inadvertently exclude a vast body of research providing key insight into AFC’s welfare impacts that falls outside standard inclusion criteria–particularly when disciplinary conventions prioritize rich qualitative reporting over standardized quantitative metrics. In contrast, a more flexible evidence synthesis could permit researchers to broadly interrogate the degree of support for positive, negative, or neutral directional associations of AFC on human and environmental well-being.

Recent advances provide new opportunities to tackle this challenge. Historically, automating environmental evidence syntheses was stymied by the heterogeneity and scale of relevant evidence. However, machine learning, transformer models, and large language models now allow researchers to increasingly automate systematic review workflows. These tools initially enhanced the efficiency of relevance screening (*36, 37*). Transformer language models (*38, 39*) led to further automation along the systematic review pipeline by robustly predicting categories of vast and diverse bodies of literature (e.g., *31, 32, 40*). But reliably categorizing nuanced information from longer texts, such as whether or not an abstract demonstrated positive impacts of AFC on subjective well-being, remained beyond reach. Now, generative pre-trained transformer (“GPT”) language models that can infer a study’s results in the context of a broader question provide the capability needed to automate such assessments (*41, 42*). Indeed, environmental science is increasingly adopting GPT models to perform sophisticated assessments of scientific evidence, ranging from extracting pairs of interacting species from abstracts to creating datasets on functional traits from species descriptions to querying large-scale data more efficiently (e.g., *41, 43, 44*). Yet, even with the rapid development of GPT models and growing experimentation in applied evidence synthesis, concerns persist about more robust methodologies that can address shortcomings such as model hallucinations and other reasoning errors (*32, 42, 45*).

Here, we report a novel approach employing multiple GPT models to overcome significant technical barriers to evaluating the co-benefits and trade-offs of AFC on human well-being and the environment. We systematically evaluated the distribution of positive, negative, and neutral associations between AFC and material human wellbeing, non-material human well-being, biodiversity, and ecosystem services ((*33, 46*), Materials and Methods). We used a standardized prompt developed through iterative expert and programmatic prompt engineering (*47, 48*). Our original sample totaled 32,273 studies predicted to contain evidence on AFC and human or environmental well-being co-impacts, and there were 97,744 distinct study and individual co-impact combinations (henceforth, “abstract”, (*33*)). For each abstract, we used five GPT models, selected based on academic and industry benchmarks, as well as a range of architectures and training data, to predict the directional impact of AFC on human material well-being, non-material well-being, ecosystem services, and biodiversity (Figure 1, Table S2, Materials and Methods). Following prior research, we applied a majority vote procedure to identify the most credible directional label for each abstract (*49*– *51*). We initially constrained our labels to positive, negative, and neutral, where neutral could indicate mixed benefits and drawbacks, no clear directional association, or that an abstract should be dropped from further consideration due to its lack of relevance to our research question. We therefore conducted a secondary screening process for neutral abstracts to exclude those abstracts that were irrelevant due to reasons including a lack of empirical information. Finally, to guard against hallucination as a mode of failure, we examined the semantic entropy of the predictions for each abstract in our final directional association dataset (n = 29,038 abstracts) (*52*).

**Figure 1:**
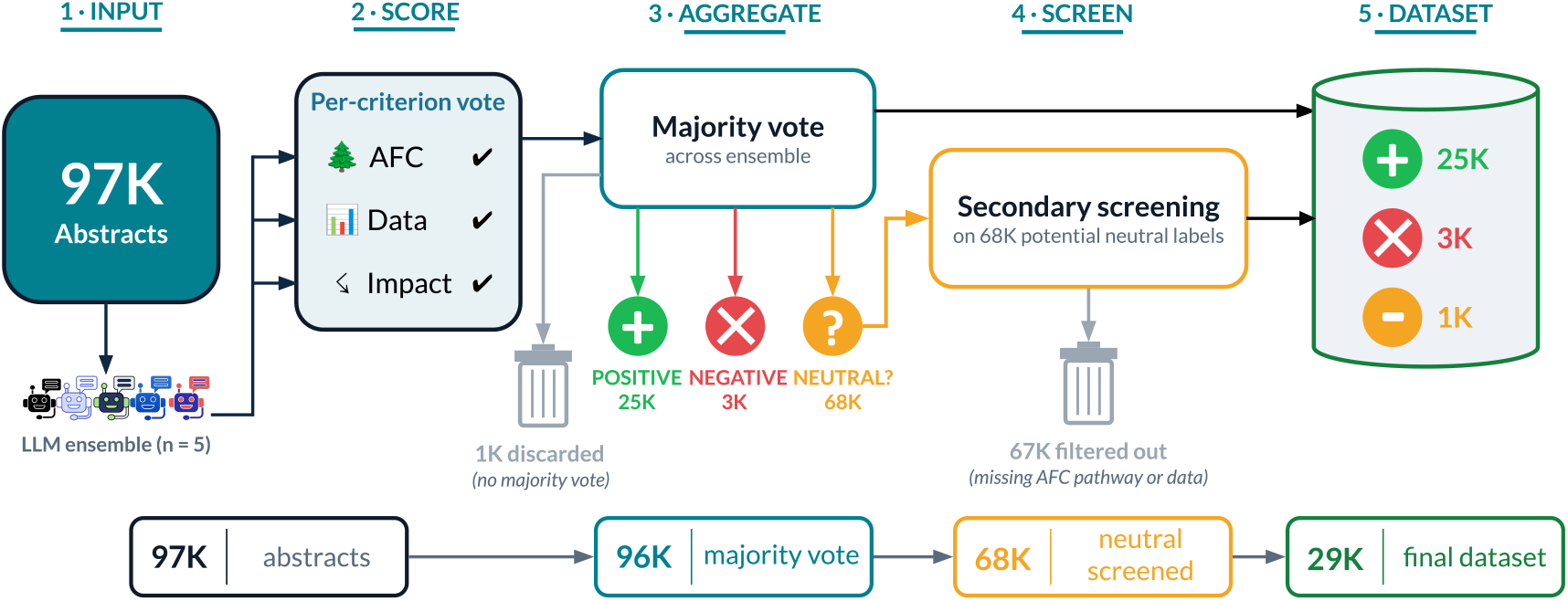
A diagram of the data processing pipeline which featured individual labels (initial set of categories: `positive, negative, neutral`) assigned by multiple GPT models, ensembling via majority vote, and secondary neutral screening to separate truly neutral abstracts from irrelevant abstracts (final set of categories: `positive, negative, neutral, irrelevant`).

Our GPT ensembling approach is analogous to using a team of coders to systematically assess evidence, and resolve disputes, for large volumes of evidence. The approach shares conceptual parallels to global synthesis efforts such as the IPBES or IPCC Global Assessment or the Lancet Commission on Health and Climate Change. These global assessments typically involve processes that seek to reach agreement among a diverse group of experts on the state of evidence rather than a formal meta-analysis that could excel at quantitative assessment at the expense of a more holistic evaluation of cumulative knowledge. Yet global synthesis efforts are still hindered by the time, effort, and expertise needed to conduct a systematic review of the evidence. Our approach presents a rigorous, replicable, and feasible pathway for assessing evidence for topics that cover a wide diversity of disciplines and an enormous volume of studies, providing a step toward the reproducible evaluation practices that are needed for GPT models to be used more rigorously in applied evidence synthesis.

We find overwhelmingly positive associations between AFC and human and environmental well-being, with AFC impacts in Africa and for material human well-being globally having the most mixed evidence. Importantly, we observed that the vast majority of the ensemble abstract labels had low semantic entropy (85% with a score of 0) and that filtering based on different thresholds did not meaningfully change our results; hence we present our findings on the full dataset. Our study indicates there is ample evidence that AFC can realize human well-being and environmental co-benefits.

## 2 Results

Across 32 years of data (1990-2022), we found that the vast majority (~85%) of the 29,038 abstracts indicated positive associations between AFC and human well-being or environmental co-impacts (Table S3), while negative (~11%) and neutral (~4%) associations were far less prevalent (Figure 2). The total volume of abstracts varied across co-impact categories, ranging from 3,256 for material human well-being to 9,984 for ecosystem services (Figure 2**A**). Collectively, all co-impacts exhibited an average annual growth rate of 12%, showing substantial research production over the three decades in our dataset. Moreover, the proportion of evidence that was positive either tended to remain steady over time (material and non-material HWB) or increased slightly (biodiversity and ecosystem services).

**Figure 2:**
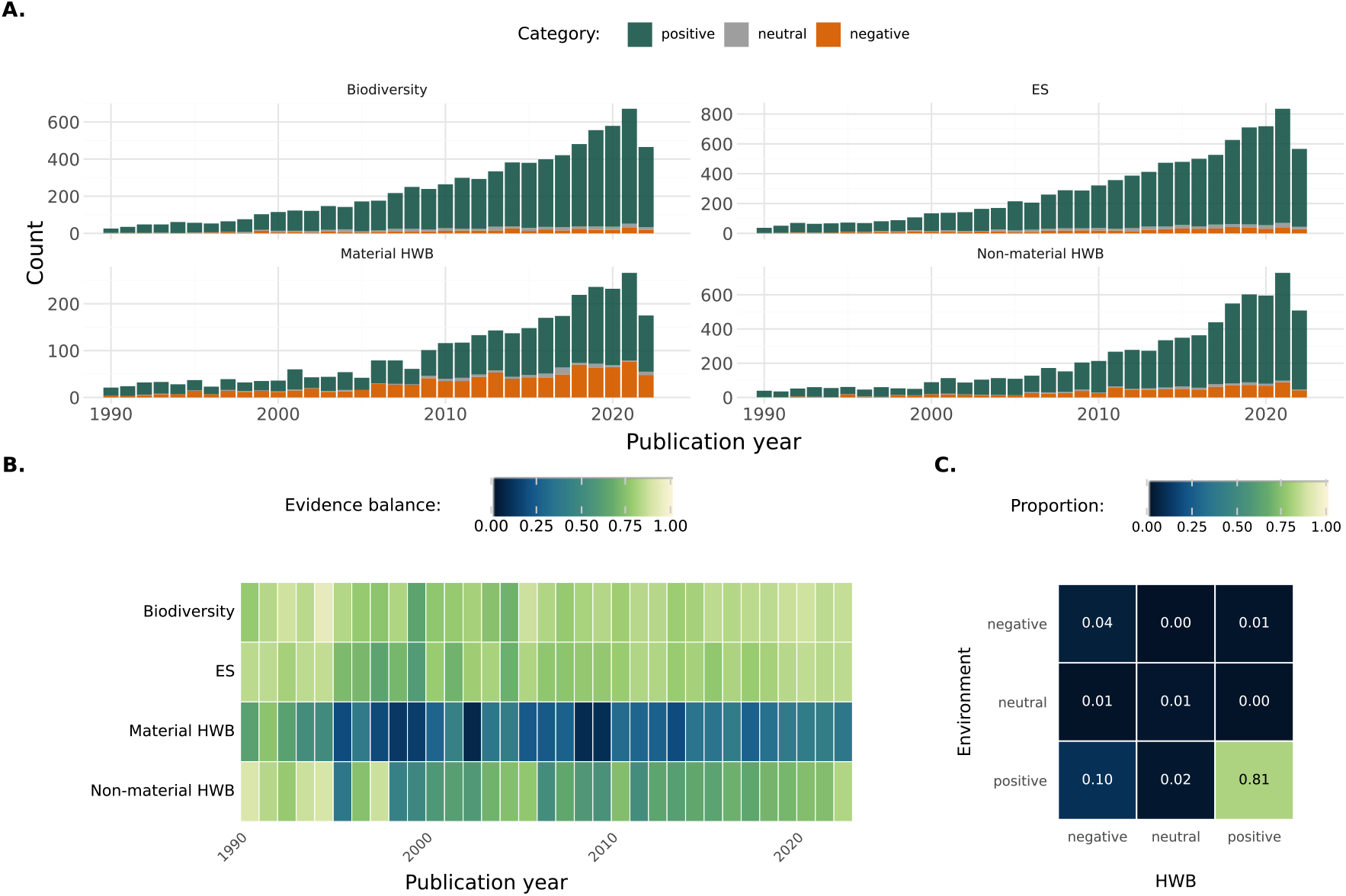
The distribution of positive, negative, and neutral associations between AFC and human non-material wellbeing, material well-being, and environmental well-being, which encompassed biodiversity and ecosystem services (**A**). Changes in the balance of evidence between positive versus all other associations, defined as *p*_*pos*_ − (1 − *p*_*pos*_) or the proportion of positive evidence minus the sum of the proportions of negative and neutral evidence (**B**). The distribution of directional associations for 3,548 abstracts containing information on at least 1 environmental (ES and/or biodiversity) and 1 HWB (Material and/or Non-Material) co-impact (**C**).

While the majority of our AFC abstracts presented positive associations, human well-being (“HWB”) dimensions– especially material HWB–tended to be less positive (Figure 2**B**). Across the thirty year span of the data, material HWB showed an upper bound of 44% negative associations and non-material HWB almost 30%. In contrast, environmental co-impacts only had a maximum of 14% negative associations with AFC. Figure 2**C** shows data on the 3,548 abstracts containing evidence for HWB and environmental co-impacts, which provide insight on potential trade-offs of AFC. These studies indicate that there are mostly synergies between human and environmental benefits (81% positive overall), but we also see trade-offs (10%) or cases where there are negative co-impacts for both people and the environment (4%). For this subset, the distribution of positive, negative, and neutral associations is largely similar to the overall sample.

There were a total of 153 countries in the 70.3% of abstracts with location data. The number of abstracts per country varied (mean: 169.5, median: 45) and ranged from 1 (e.g., Uzbekistan, Martinique) to several thousand (e.g.,

Mexico, China, Indonesia, Brazil, USA), mirroring geographic bias commonly found in environmental studies (*53*). At the country scale, similar to the aggregate trends, HWB tended to exhibit less positive evidence than biodiversity and ecosystem services (Figure 3; mean evidence balance for HWB of 67% versus 85% for the environment, Table S4). Some countries exhibited win-win patterns for humans and the environment from AFC, such as Brazil, China, Colombia, Ethiopia, or Kenya. Others showed trade-offs, where the evidence suggested greater benefits for the environment than HWB (e.g., Indonesia, D.R. Congo, Peru) or for HWB rather than the environment (e.g., South

**Figure 3:**
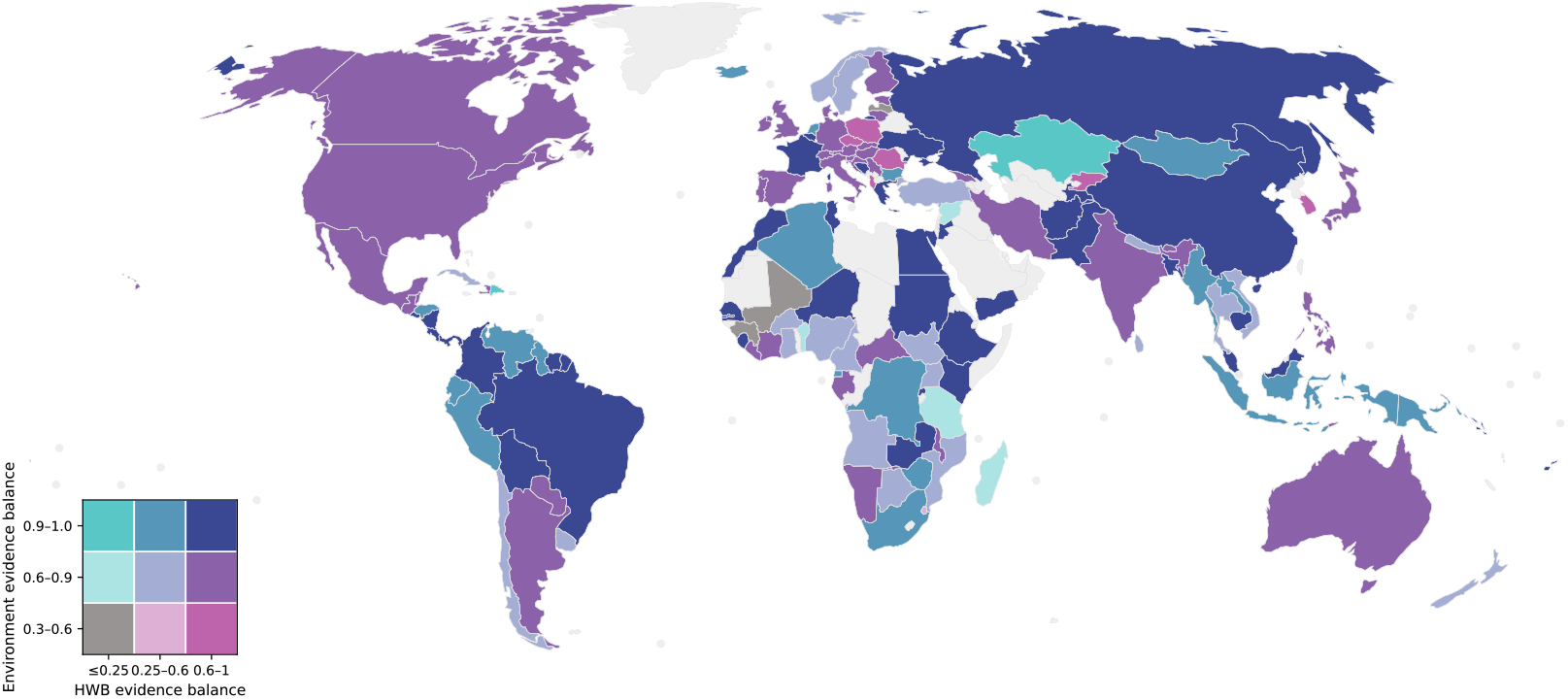
The evidence balance (*p*_*pos*_ − (1 −*p*_*pos*_)) values for human well-being versus environmental co-impacts across countries. We required that each country have at least 5 abstracts to be mapped, dropping our sample from 153 countries to 131.

Korea, Poland, Romania), though the latter was a much smaller category. Few countries had relatively negative evidence overall for all categories of AFC co-impacts (e.g., Mali where *p*_*pos,env*_ = 0.67 whereas *p*_*pos,HWB*_ = 0 or Guinea where *p*_*pos,env*_ = 0.71, *p*_*pos,HWB*_ = 0.6). Overall, there were more countries with more highly positive evidence (defined as *p*_*pos*_ *≥*0.8 *⇒* evidence balance *≥*0.6) for environmental co-impacts than HWB (HWB = 88 countries with highly positive evidence out of the 131 mapped countries, environment = 122).

Crosswalking our evidence map with data on past deforestation and future conversion pressure (*54*–*56*), we identified quadrants of forest conversion risk (Figure 4). The top left and right quadrants highlight countries of interest for future conservation action: *Frontline hotspots*, which exhibited relatively high past conversion rates and future conversion risks (e.g., Cambodia, Uganda, Honduras), and *Emerging frontiers* where projected future conversion risks are high and past deforestation rates were low (e.g., Benin, Ecuador, Sri Lanka). We observed that countries with the greatest future conversion risk tended to have less evidence overall (Figure 4, Table S5); countries with the most evidence generally exhibited lower average future conversion pressure on their forests (mean ± *SE*, all countries = 238.8 ±46.5, low future threat countries = 342.8 ±96.4, *Emerging frontiers* = 126.5 ±32.4, *Frontline hotspots* = 162.8 ±34.7). These frontier areas represent critical locations where evidence is most urgent given their vulnerability to forest conversion.

**Figure 4:**
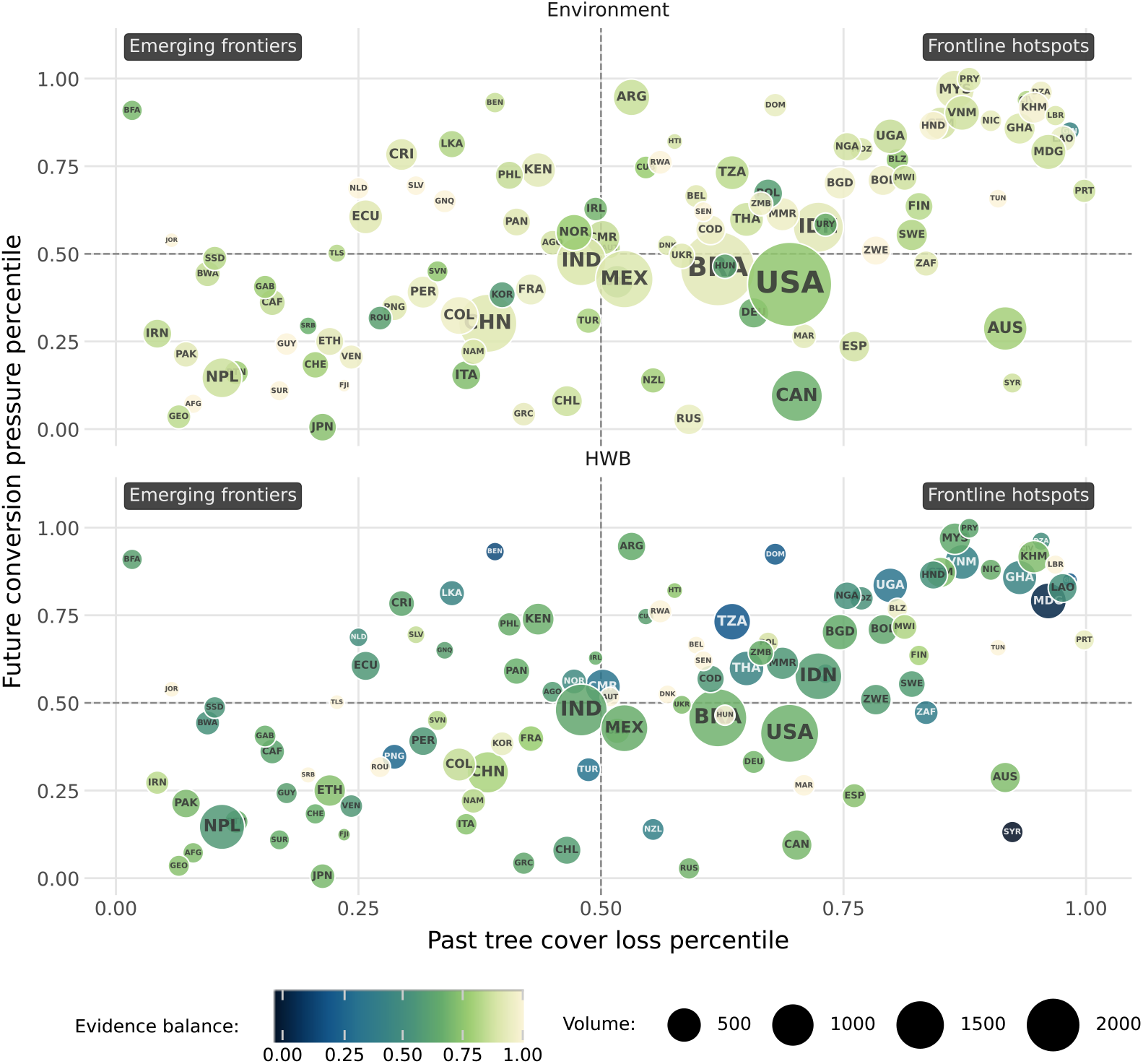
The distribution of AFC evidence given past (*54*) versus future conversion rates (*55*) for natural forests globally (*56*). Countries depicted in the figure had at least 5 studies for each category of co-impact (Environment and HWB, totaling a minimum of 10 studies), which resulted in a final sample size of 107 countries.

Sparse evidence is particularly concerning when more people may experience the consequences of forest loss or conservation, based on evidence-gap frameworks that account for population exposure or the potentially impacted population for climate impacts (*57* –*59*). Our approach for characterizing sparse evidence is to calculate a *research attention* metric based on the number of publications per forest-proximate people (“FPP”, Newton *et al*. (*21, 22*)). We do so for *Emerging frontier* and *Frontline hotspot* countries. Low values (those below the median, which was 1 abstract per 150,000 FPP) are flagged as limited *research attention* relative to the size of the potentially affected population. We then combined our *research attention* metric with data on mitigation potential (Figure S3, Table S5), and observed concerning gaps for countries with high deforestation risk, high FPP, and substantial mitigation potential, such as the Philippines, Nigeria, Rwanda, Myanmar, or the Democratic Republic of the Congo. This sparsity of evidence encompasses 354*M* forest-proximate people in future deforestation hotspots on the African continent, 258*M* FPP in greater Asia, 45*M* FPP in the European continent, and 19.6*M* FPP in the Americas.

### 2.1 Robustness check for publication bias

Publication bias threatens the validity of any evidence synthesis. We evaluate this to the best of our ability using our dataset’s metadata and external data, where publication bias may be apparent if the distribution of positive, negative, and neutral associations differ across observable factors. We find broad consistency in the distribution of positive, negative, and neutral AFC impacts across tiers of journal selectivity and the scientific attention received by a study, proxied by its citation count (Figures S4, S5). Given our observed disparities in AFC publication rates, we sought to examine if more equitable representation across different global cultures or income levels would shift the evidence map of AFC impacts. Our analysis reveals no significant impact of continent on the degree of positive AFC evidence, as a proxy for different global cultures, and contrary to initial expectations, we saw some suggestion that lower-income regions might actually lead to an even larger degree of positive evidence for AFC co-benefits (Table S6). However, publication bias can reflect the choices that authors make that are unobservable to the outside world; for example, an author confronted with unexpected or non-significant findings may choose not to write up their results. This potential driver is, however, beyond the scope of what we can reasonably assess.

## 3 Discussion

AFC has long been central to environmental conservation and nature-based climate solutions (*1, 3, 6, 10*). Our results reiterate that, globally, AFC indeed provides substantial benefits for local communities and the environment. Across a broad set of well-being dimensions, we offer important empirical support for what were previously presumed cobenefits of AFC for people and nature. Nevertheless, our results caution that in particular regions, there may be trade-offs for AFC and human well-being, and in particular, material human well-being. There is a rich literature on how AFC policies often fail to benefit local communities, creating politically fragile policies that in the end may fail to achieve conservation and broader human development goals. More recently, the momentum for 30×30 targets has been accompanied by growing calls to carefully and explicitly consider trade-offs of AFC policies (*7, 19*). Our study overcomes significant technical and resource barriers, presenting a global and comprehensive assessment on the current state of evidence of AFC and key co-impact categories.

Although we observe that AFC should generally lead to positive impacts for both material and non-material human well-being, there are countries where the evidence is sparse or suggestive of potential trade-offs resulting from AFC. For the roughly 677*M* forest-proximate people in countries with scant evidence, addressing these knowledge gaps is critical for more effective and equitable AFC implementation. Limited evidence for *Emerging frontier* and *Frontline hotspot* countries is also concerning. This is especially true for countries with large populations of forest-proximate peoples such as Nigeria, the Philippines, or Côte d’Ivoire. Given that local community support and shared benefits are critical for durable forest protection (*18, 20, 27, 28*), having a stronger base of evidence to guide AFC interventions in these regions is of critical importance.

There has long been an assumption that AFC inherently prioritizes conservation goals over local livelihood opportunities. This premise has driven substantial research efforts in benefit-sharing, focusing on how instruments such as payments for ecosystem services are needed to offset real or perceived AFC trade-offs for frontline and fence-line communities (*60, 61*). Our results show that the majority of literature reports that AFC is capable of delivering HWB benefits. The combination of our results and the persistence of deforestation suggest deeper societal factors driving land conversion and obstacles to benefit-sharing. Institutional failures can prevent community co-benefits (*17, 27, 28*), and can be compounded by forces such as large-scale land acquisitions where profits are captured far above the local level (*62*). The distribution of positive AFC impacts for HWB and the environment highlights that the mechanisms and factors moderating benefit flows to communities continues to be an urgent research priority; effective conservation requires moving beyond localized incentives toward larger-scale governance deficits, as our results point to the importance of identifying and understanding institutional failures at regional, national, and international levels alongside any local incentives to protect or clear forests.

Our GPT ensembling approach, alongside using semantic entropy (*52*) to test for substantial hallucination-induced reasoning errors, presents an advance in employing the latest AI tools to evidence synthesis. Continued methodological advances offer exciting possibilities for dynamically tracking and quantifying vast bodies of evidence for global priorities. While there are many concerns around ethics, data sovereignty, privacy, and the environmental footprint of using these models (*63, 64*), recent work, and our results here, show a path forward with an ensemble of models, which can include smaller, open-weight models (*51, 52, 65, 66*). Emerging work shows that smaller language models and open-source platforms can robustly power retrieval-augmented generation systems where the public can ask nuanced questions of academic evidence (*67* –*69*). These tools are rapidly evolving, and we believe a priority for applying these tools to evidence synthesis is to generate workflows that are transparent and replicable.

Nevertheless, while research on scientific agent workflows for question-answering, hypothesis, or code generation is growing (*41, 67, 69*), AI for open scholarship should also prioritize the type of robust structured information extraction that would permit for more scalable meta-analyses (*70*–*72*). Automating meta-analyses of full-length scholarly articles remains a challenge, especially in highly interdisciplinary fields such as environmental science and studies (*73*); researchers may publish in thousands of venues and across hundreds of disciplines, each of which has their own norms and practices for conveying findings. Research indicates that generating reliable meta-analytic data is stymied by issues with data ingestion and quality, and that even frontier large language models struggle to produce the correct number of results and results that are correctly linked with their context (e.g., a study, a population, an intervention or control condition, and any other qualifiers needed to make sense of a measurement or descriptive result, (*70, 72, 73*)). Strong results for structured information extraction for inquiries with focused disciplinary constraints, often limited to shorter spans of input data (e.g. abstracts or clinical trial reports), highlight what could be possible for deeper and broader studies aiming to compass full-length scholarly articles (*71*).

Our work provides support that AFC, as an immediate biodiversity and climate solution, is poised to offer benefits for local human communities and advance conservation outcomes. We illustrate a path forward for robust, large-scale, and rapid syntheses of multidisciplinary evidence bases. More broadly, our approach can be applied across any intervention if the evidence base is identified and requisite data exist, ranging from restoration, management of working lands, illegal wildlife trafficking, to sustainable supply chains. More specifically to AFC, we believe future research efforts can build on our results and continue to leverage technical advancements to assess the size, sustainability, timing, or other features of AFC co-impacts that are salient for implementation. More nuanced assessments of what types and elements of AFC interventions lead to co-benefits rather than trade-offs in specific geographies will provide insights that can directly inform regional and local conservation and climate strategies. AFC continues to be integral to contemporary conservation and sustainable development efforts, and a critical next step will be to go beyond a general understanding that positive impacts are possible to support fair and equitable implementation efforts that maximize HWB and environmental co-benefits.

## 4 Materials and methods

### 4.1 An initial evidence map of conservation and climate mitigation pathways

Chang *et al*. (*33*) used transformer language models to map 257,266 studies to different ecosystem-based intervention pathways and co-impacts to people and nature (*2, 46*). These pathways spanned protection, management, and restoration actions (*2*), while the co-impacts encompassed human material and non-material well-being, biodiversity, and ecosystem services ((*33*), Table S1). Ultimately, this dataset yielded 32,273 studies associated with avoided forest conversion (“AFC”). As each abstract could have multiple co-impacts, our final sample was 97,744 abstract and co-impact combinations (henceforth: “abstract”).

### 4.2 Assessing the directionality of avoided forest conversion studies

We took the majority vote of a set of generative pre-trained transformer (“GPT”) large language models to label the 97,744 abstracts (Figure 1). In the prompt for all models, we evaluated whether there was a clear directional association between AFC and impacts to people and nature (Figure S1). We developed our prompt through a combination of expert iteration and programmatic prompt engineering (*47, 74, 75*). At the end of the prompt, the model was asked to return one of three categories for an abstract with respect to AFC and its impacts on people or the environment: `positive, negative,` or `neutral`. In this initial screen, `neutral` could indicate mixed benefits and drawbacks (true neutrality) or no clear directional association (irrelevance). We used prompting features that have been shown to improve in-context learning for more accurate generation: chain-of-thought prompting (*65, 76*), step-by-step reasoning about the evidence contained in an abstract (*48*), and reinforcing the abstract’s context through text summarization to arrive at a more grounded output (*49, 77*) (for more details, please consult the SI).

For all abstracts initially labeled neutral (n=68,707), we used a smaller GPT model (OpenAI `gpt-5-mini-2025-08-07`) for a secondary inspection, based on research examining how the output of larger models can serve as robust and informative inputs for smaller models (*66, 78, 79*). This second-stage check occurred only when 1) the ensemble majority vote decision was neutral, and 2) the prediction of an individual GPT model was neutral. The check examined whether the abstract contained clear evidence for neutrality. Abstracts that lacked empirical evidence or had insufficient and unclear information were `irrelevant` and excluded. These labels were then combined with the original directionality outputs to produce a final four-class prediction (`positive, negative, neutral`, or `irrelevant`). Approximately 1% of the abstracts had no clear majority vote (result: tie, n=1,049); these were excluded from additional analyses.

Prior work has shown that ensembling model predictions, and in particular, a simple majority-vote aggregation, can lead to more reliable results from a GPT data-labeling pipeline (*49*–*51*). We chose GPT models based on academic benchmarks (*41, 80*) and industry evaluations of model performance (e.g., HuggingFace Open LLM Leaderboard and ArtificialAnalysis.ai), aiming to capture a diversity of architecture, training regimes, and reasoning performance. An ablation study of different ensemble sizes (ranging from the single best model to an ensemble of 7 models) demonstrated that 5 models (GPT-5, Gemini 2.5 Pro, Qwen3-235B, Llama 3.1, DeepSeek Reasoner) yielded the best performance on our data (Table S2).

For each abstract, we collected all predictions, one per model (n=5 per abstract), and computed the most frequent label from the final four class prediction set of `positive, negative, neutral,` and exclude. We evaluated these results against a randomly selected sample of 100 abstracts that were independently coded by four members of the author team who have expertise in biodiversity conservation, nature-based climate change mitigation, and human well-being and ecosystem service co-benefits. Our ground truth label was the majority vote of the four human coders. We computed accuracy and weighted-average *F*_1_ scores for the ensembled prediction. We found that ensembling substantially improved reliability relative to any single model, but that gains diminished beyond a moderate number of models. In particular, we found that an ensemble of 5 models (OpenAI GPT 5, Gemini 2.5 Pro, Qwen3 235B, Llama 3.1, Deepseek R1) consistently achieved the strongest balance of accuracy and *F*_1_ scores (Table S2); larger ensembles introduced additional disagreement without corresponding performance improvements. The overall performance of the ensembled GPT predictions was comparable to or better than the expert human coders (expert range: [0.67, 0.76], best GPT ensemble: 0.78, Table S2), and within range of recent research for more nuanced data coding using large language models (*43, 57, 58*). Based on our review of these data, we concluded that the discrepancies between GPT and expert labels reflected intrinsic ambiguity in the abstracts that was even challenging for human experts to interpret, rather than methodological issues such as systematic model parametric memory issues or reasoning errors.

### 4.3 Robustness check with semantic entropy

To control for potential LLM data coding reliability issues, we used a recent method that leverages the uncertainty across multiple LLM outputs. Specifically, we calculated the semantic entropy of each abstract’s reasoning outputs across the five GPT models, which detect for confabulations, a type of LLM hallucination characterized by being both arbitrary and incorrect (*52*). This method clusters LLM outputs based on whether the responses are consistent with one another, or contradictory. The larger the number of contradictory, and thus distinct, reasoning responses, the greater the number of clusters, and vice versa. A larger number of clusters corresponds to greater uncertainty.

The range of observed entropy values for the abstracts in our dataset was [0, 1.05], corresponding to [1, 3] distinct reasoning clusters. We observed that the vast majority of the ensemble abstract labels had low entropy (85% with a score of 0). Ultimately, the overall sample size, distribution of values, and proportion of `positive, negative`, and `neutral` labels were relatively consistent given different semantic entropy filtering thresholds (Figure S2). Given our observation that filtering would not substantively change our results, our analyses used the full dataset.

### 4.4 Analyses

We cross-walked our AFC evidence with additional datasets to examine geographic patterns in past versus future forest conversion threat, as well as forest stock versus drivers of forest dependence. We detected location data in 20,410 abstracts and aggregated these data to 153 sovereign entities (henceforth, countries). We used observed rates of country-level tree cover loss (*54*) and the Conversion Pressure Index (“CPI”, (*55*)) masked to pixels predicted to be natural forest (“SBTN”, (*56*)) as proxies for past and future forest conversion, respectively. As the CPI and SBTN rasters exhibited different resolutions (1×1km versus 30×30m), we first resampled the coarser CPI raster to a finer resolution matching the SBTN raster, and then masked the CPI raster to only those pixels containing natural forest. We then aggregated the masked CPI pixels within country-level polygons and calculated the mean, median, and number of pixels. Above-ground biomass at country scale, measured as average tonnes per hectare, served as a proxy for forest stock. As a measure of forest dependence, we used the proportion of forest-dependent people within a 5 km radius (*22*). All analyses were conducted in Python (version 3.12).

## Acknowledgments

We thank Nikhil R. Deshmukh, Amrita Gupta, and others for thoughtful feedback that improved the project. We acknowledge funding support from the Bezos Earth Fund and other donors supporting The Nature Conservancy. The authors declare no competing interests.

## Notes

### Competing Interest Statement

The authors have declared no competing interest.

