## Supplementary Information for "A global assessment reveals widespread co-benefits for forest protection"

### 1.1 Compiling studies on Avoided Forest Conversion

Chang *et al.* (*1*) defined Avoided Forest Conversion ("AFC") following Griscom *et al.* (*2*) and Nature4Climate 2023. This definition specifies that AFC is "Avoided forest conversion, disturbance, degradation, fragmentation, and deforestation of any kind of forest (i.e., rainforests) and natural woodland, including within and close to urban areas." Co-impacts were defined based on McKinnon *et al.* (*3*) and Pörtner *et al.* (*4*), and encompassed 9 categories which we grouped into 4 broader categories (Table S1). Two of the 9 co-impact categories occurred at extremely low frequencies in our evidence mapping dataset, and thus, we excluded these from additional analyses. For more information on the initial sampling strategy, please refer to Tables S9-S14 in Chang *et al.* (*1*).

| Co-impact | Count | Category |
|---|---|---|
| Economic or material living standards | 3,256 | Material HWB |
| Security and safety | 2,796 | Non-material HWB |
| Subjective well-being | 2,075 | Non-material HWB |
| Culture and spirituality | 1,467 | Non-material HWB |
| Health | 1,151 | Non-material HWB |
| Ecosystem services | 9,984 | Ecosystem services |
| Biodiversity conservation | 8,149 | Biodiversity |
| Social relations | 127 | Excluded, due to a much smaller and potentially non-representative sample size |
| Education | 33 | Excluded, due to a much smaller and potentially non-representative sample size |

Table S1: Definition and grouping of the co-impacts of AFC

### 1.2 Prompt development, data labeling process, and semantic entropy checks

An effective prompt presents a task for a GPT model that resembles the types of tasks that the model originally encountered in training (e.g. next word prediction; (*5*, *6*)). Prompts can be used for zero-shot (generate output based on the input string alone) or few-shot (provide several solved examples of the new task in the input string) in-context learning (*6*, *7*). We developed a prompt for our data coding task through an iterative, expert-led process that was complemented with programmatic prompt engineering (*8*). To produce a more consistent output format, we used `DSPy` to traverse embedding space and optimize components of the prompt given an objective (*8*). In our case, the objective was generating correctly labeled output in the format of {`impact: 'biodiversity', direction: 'positive'`}.

Our labeling task entailed several challenges for GPT as a data coder: inductive reasoning (determining whether or not particular features, such as forest conversion, were present in an abstract), combining multiple inferred outputs, and a substantial number of input tokens (*7*, *9*, *10*). To address these challenges, we developed a prompt that featured chain-of-thought prompting (*11*, *12*), step-by-step reasoning about forest conversion and impacts to people or nature (*7*, *11*, *13*, *14*), and reinforcing the abstract context in the prompt through a summarization process in the reasoning chain (*15*–*20*) before producing the final response output. Figure S1 displays the final prompt.

Our prompt development process identified several key features: 1) focusing on forest conversion rather than avoided forest conversion, as the latter tended to induce more substantial reasoning errors, and 2) excluding an "irrelevant" category in the primary screening process as that tended to promote "lazy" or "shortcut" reasoning errors where `neutral` labels were too often and inconsistently assigned to other categories (*9*, *21*). We therefore added a secondary neutral versus irrelevant screening stage in our process. This was informed by a manual inspection of 1000 neutral abstracts which confirmed two broad categories: 1) abstracts that lacked empirical data and were thus irrelevant, versus 2) abstracts that displayed a truly neutral association between AFC and a co-impact, often due to mixed positive and negative evidence (e.g., community access to firewood decreased but surrounding community incomes increased). In conducting the secondary neutral screening process, we used a smaller model. Our choice was inspired by research on multiple GPT models inspecting output traces (*12*, *19*, *22*, *23*), which has shown that larger models and their chain-of-thought rationale outputs can act as instructional inputs to smaller models (*17*, *24*, *25*).

Table S2 presents the results of comparing different GPT model combinations (ensembles of models) on coding AFC abstracts. The starting set of models was selected based on prior research (*26*, *27*) and industry benchmarks (*28*, *29*). Each ensemble's performance was compared to a set of 100 abstracts that were quadruply coded by four experts in biodiversity conservation. We observed that in the case of the best ensemble ($n = 5$), the majority GPT vote categorization exceeded some of the individual human coders' performance versus the majority expert vote.

| **Ensemble (best composition at each size)** | **Accuracy** | **Weighted $F_1$** |
|---|---|---|
| 1 model: GPT-5 | 0.76 | 0.78 |
| 2 models: GPT-5, GLM-4.6 | 0.76 | 0.78 |
| 3 models: GPT-5, Claude Sonnet 4.5, Qwen3-235B | 0.77 | 0.78 |
| 4 models: GPT-5, Gemini 2.5 Pro, Llama 3.1, DeepSeek Reasoner | 0.77 | 0.79 |
| **5 models: GPT-5, Gemini 2.5 Pro, Qwen3-235B, Llama 3.1, DeepSeek Reasoner** | **0.78** | **0.79** |
| 6 models: GPT-5, Claude Sonnet 4.5, Gemini 2.5 Pro, Qwen3-235B, Llama 3.1, DeepSeek Reasoner | 0.77 | 0.78 |
| 7 models: GPT-5, GLM-4.6, Claude Sonnet 4.5, Gemini 2.5 Pro, Llama 3.1, Qwen3-235B, DeepSeek Reasoner | 0.75 | 0.76 |

Table S2: Performance of the best-performing model ensemble at each ensemble size on the $N = 100$ human-coded validation set where the ground truth was the majority vote of four expert human coders. A 5-model ensemble achieved the strongest balance of accuracy and weighted $F_1$, while larger ensembles produced reduced performance and agreement.

Figure S2 displays the impact of using semantic entropy to filter the dataset. We observed that the majority of the AFC abstracts exhibited low entropy and that there would be no substantive change to the overall distribution or proportion of positive, negative, and neutral abstracts.

Table S3 displays the directional associations for each category of co-impact across the totality of the dataset. These results show that the majority of the evidence demonstrates positive associations with AFC.

| Co-impact category | Overall directional evidence | Proportion |
|---|---|---|
| Biodiversity | positive | 0.91 |
| | negative | 0.05 |
| | neutral | 0.04 |
| Ecosystem services | positive | 0.89 |
| | negative | 0.06 |
| | neutral | 0.04 |
| Material HWB | positive | 0.66 |
| | negative | 0.31 |
| | neutral | 0.03 |
| Non-material HWB | positive | 0.84 |
| | negative | 0.14 |
| | neutral | 0.02 |

Table S3: The overall directional evidence for each co-impact category, presented as proportions.

A Priming the LLM to act with expert knowledge

B Defining task outcomes: Here we specify the labels for the "directional association" zero-shot reasoning task

### System prompt

You are an environmental science expert reviewing scientific abstracts to compile evidence for the impacts, positive, negative, or neutral, of forest conversion on ecosystem services.

Your job is to determine the directional association (Positive, Negative, or Neutral) between forest conversion and ecosystem services based on empirical evidence provided in the abstract.

C Definitions: We prime the LLM to draw on certain areas of indexed knowledge by defining key terms

D Ruleout conditions: We inform the LLM about how to deal with categories that are not clearly one or the other

E Reverse logic: Instruct the LLM to differentiate between forest conversion versus protection and introduce the "reverse logic" sub-task

F Constrain reverse logic: Here, we instruct GPT on what categories constitute forest conversion versus its opposite

### Background information

Keep in mind:

- forest conversion refers to typically human-induced disturbance, degradation, fragmentation, and deforestation of any kind of native forest and natural woodland, including within and close to urban areas.

- the directional association between forest conversion and ecosystem services is Positive if converting the forest through the above mentioned activities would overall benefit ecosystem services.

- the directional association between forest conversion and ecosystem services is Negative if converting the forest through the above mentioned activities would overall be disadvantageous for ecosystem services.

- the directional association between forest conversion and ecosystem services is Neutral if it is not conclusively Positive and not conclusively Negative.

- those outcomes of forest conversion, which only sustain, maintain, or do not affect native forest-level attributes are neither Positive nor Negative.

- if the abstract focuses on the outcome(s) of protection or restoration of a forest habitat, then instead of forest conversion, we call that process avoided forest conversion. The directional association between forest conversion and ecosystem services is always the opposite of the directional association between avoided forest conversion and ecosystem services.

- compared to native forest or natural woodland, forestry management practices such as reduced impact logging or extended rotations in uneven-aged stands are not avoided forest conversion practices, but are instead a form of forest conversion where the objective is to exert less negative impact on the forest landscape compared to business-as-usual practices.

- on the other hand, management practices such as community-managed forests or joint forest management or other practices that seek to protect, restore, or rehabilitate native forests and natural woodlands threatened by commodity or shifting agriculture, urbanization, or intensive logging can constitute avoided forest conversion.

- given the two points above, it is critical to carefully consider the types of forest management, if any, discussed in the abstract to carefully determine which elements in the abstract correspond to forest conversion.

G Chain of thought prompting

H Text extraction task: ensure that the LLM references and draws on the language in the abstract

I Text summarization helps GPT arrive at a more principled decision that reduces the risk of hallucination

J Ensuring that the output follows the desired format

### Solving the task

In order to solve the task step by step, extract the following information from the abstract:

- What embodies ecosystem services in the given abstract?

- Identify the forest habitat under study, even if it has to be inferred from the abstract text.

- Identify the type of habitat(s) used for comparison, if any, even if they have to be inferred from the abstract text. They are often the studied forest habitat but under different conditions (e.g. in case the native forests go through the above mentioned human-induced activities).

- Identify which of the forest habitats are the native forests and which are the converted forests?

- Identify empirical data presented in the abstract that illustrates a comparative analysis regarding ecosystem services within the native and converted forest habitats in question.

- Identify the direct impacts attributed to forest conversion (i.e. moving from native forests toward converted forests) concerning ecosystem services.

- Examine whether or not the abstract focuses on the outcome(s) or impact(s) of restoration, rehabilitation, or protection of a forest habitat. If yes, then the abstract discusses the impact(s) of avoided forest conversion, and therefore reverse logic must be applied in order to report the impact(s) of forest conversion.

- Now summarize what we know at this point and make your decision regarding directional association, based on the impact that forest conversion has toward ecosystem services. Do not speculate and do not consider general outcomes, only the specific ones discussed in the abstract related to ecosystem services!

---

Follow the following format.

Abstract: ${abstract}

Reasoning: Let's think step by step in order to ${produce the directional_association}. We ...

Directional Association: Neutral, Positive or Negative

---

Figure S1: The engineered prompt used to process avoided forest conversion abstracts along with annotations describing elements of the prompting process.

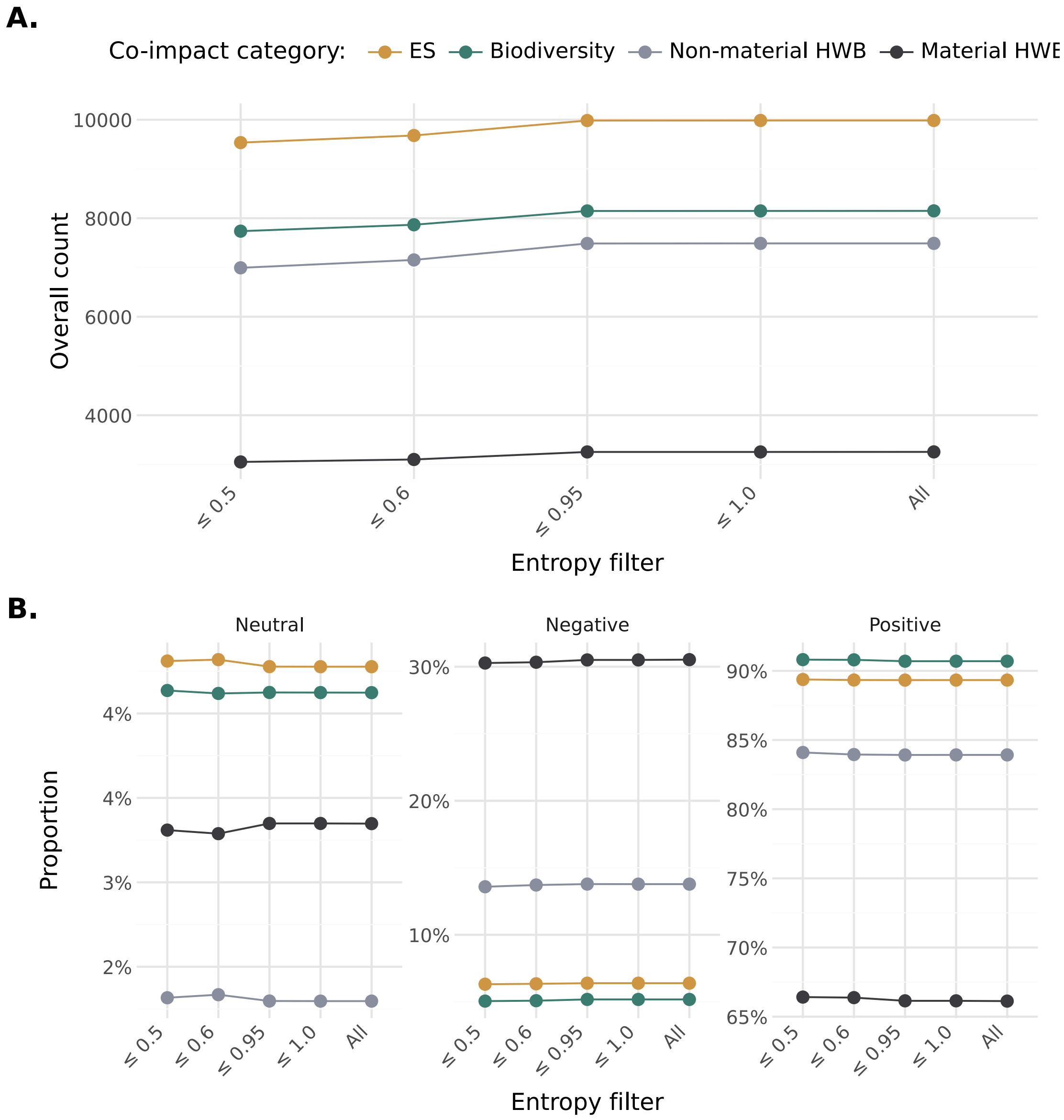


Figure S2: Using semantic entropy to conduct a robustness check, we display how the overall sample size (**A**) and proportion of positive, negative, and neutral abstracts (**B**) would change under different entropy filtering thresholds.

### 1.3 Country-level results

Table S4 displays the overall quantity of evidence (Total) and the proportional breakdown of positive, negative, and neutral evidence for all of the countries in the dataset. We collapse the two human well-being (Material and Non-material HWB) and two environmental co-impacts (ES and Biodiversity) into the categories HWB and Environment, respectively.

Table S5 shows which *Emerging frontier* or *Frontline hotspot* countries have different data coverage and carbon mitigation potential. Carbon mitigation potential is proxied by aboveground biomass (AGB), and binned into one of three categories (Low, Med, High) based on the rank of a country's AGB versus all other countries globally.

Figure S3 displays a treemap of forest-proximate populations ("FPP") and aboveground mitigation potential ("AGB"), with shading delineating the volume of AFC co-impact evidence (*research attention*: Data gap or Mapped categories).

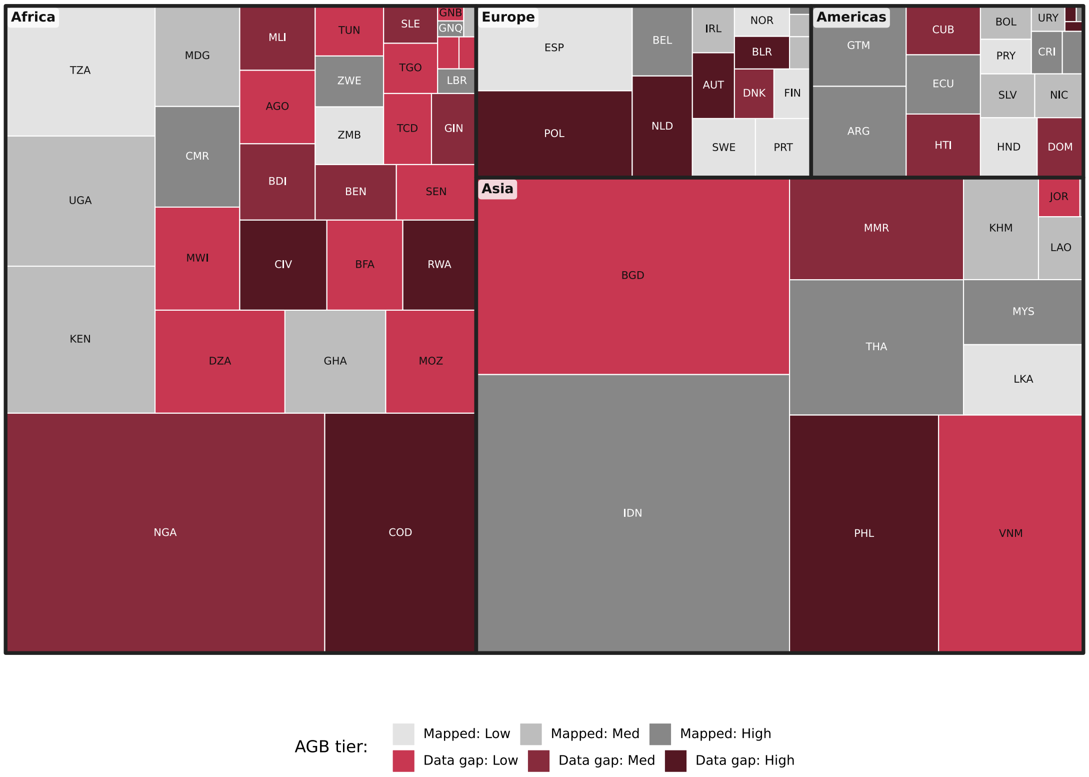


Figure S3: The distribution of *research attention* versus aboveground biomass (AGB). The sizing is based on forest-proximate populations ("FPP", (*30*)), scaled according to their proportional contribution to each continent's overall FPP, and shaded according to mitigation potential (proxied by aboveground biomass, (*31*)). Countries in red are a priority based on our *research attention* metric: they are either missing from the evidence base entirely or have a relative paucity of evidence (Data gap). Gray countries, in contrast, are those that have relatively abundant *research attention* (Mapped).

Table S4: The distribution of positive, negative, and neutral evidence for all of the countries in the dataset, divided by either human well-being (HWB) or environmental co-impacts.

| ISO3 | Name | Domain | Total | $p_{pos}$ | $p_{neg}$ | $p_{neu}$ |
|---|---|---|---|---|---|---|
| AFG | Afghanistan | Environment | 10 | 1.00 | 0.00 | 0.00 |

| | | | | | | |
|---|---|---|---|---|---|---|
| AFG | Afghanistan | HWB | 14 | 0.86 | 0.14 | 0.00 |
| AGO | Angola | Environment | 32 | 0.91 | 0.03 | 0.06 |
| AGO | Angola | HWB | 12 | 0.67 | 0.08 | 0.25 |
| ALB | Albania | Environment | 10 | 0.70 | 0.30 | 0.00 |
| ALB | Albania | HWB | 1 | 1.00 | 0.00 | 0.00 |
| ARG | Argentina | Environment | 204 | 0.91 | 0.05 | 0.04 |
| ARG | Argentina | HWB | 74 | 0.82 | 0.18 | 0.00 |
| AUS | Australia | Environment | 416 | 0.86 | 0.08 | 0.06 |
| AUS | Australia | HWB | 110 | 0.85 | 0.13 | 0.03 |
| AUT | Austria | Environment | 17 | 0.88 | 0.06 | 0.06 |
| AUT | Austria | HWB | 12 | 1.00 | 0.00 | 0.00 |
| AZE | Azerbaijan | Environment | 4 | 1.00 | 0.00 | 0.00 |
| BDI | Burundi | Environment | 1 | 1.00 | 0.00 | 0.00 |
| BDI | Burundi | HWB | 2 | 1.00 | 0.00 | 0.00 |
| BEL | Belgium | Environment | 44 | 0.95 | 0.05 | 0.00 |
| BEL | Belgium | HWB | 9 | 1.00 | 0.00 | 0.00 |
| BEN | Benin | Environment | 14 | 0.93 | 0.07 | 0.00 |
| BEN | Benin | HWB | 7 | 0.57 | 0.43 | 0.00 |
| BFA | Burkina Faso | Environment | 20 | 0.80 | 0.10 | 0.10 |
| BFA | Burkina Faso | HWB | 20 | 0.80 | 0.20 | 0.00 |
| BGD | Bangladesh | Environment | 114 | 0.97 | 0.01 | 0.02 |
| BGD | Bangladesh | HWB | 175 | 0.82 | 0.17 | 0.02 |
| BGR | Bulgaria | Environment | 6 | 1.00 | 0.00 | 0.00 |
| BGR | Bulgaria | HWB | 3 | 0.67 | 0.33 | 0.00 |
| BIH | Bosnia and Herzegovina | Environment | 6 | 1.00 | 0.00 | 0.00 |
| BIH | Bosnia and Herzegovina | HWB | 3 | 1.00 | 0.00 | 0.00 |
| BLR | Belarus | Environment | 5 | 0.60 | 0.00 | 0.40 |
| BLZ | Belize | Environment | 36 | 0.86 | 0.11 | 0.03 |
| BLZ | Belize | HWB | 22 | 0.95 | 0.00 | 0.05 |
| BOL | Bolivia | Environment | 115 | 0.97 | 0.02 | 0.01 |
| BOL | Bolivia | HWB | 108 | 0.81 | 0.18 | 0.01 |
| BRA | Brazil | Environment | 1779 | 0.94 | 0.03 | 0.03 |
| BRA | Brazil | HWB | 930 | 0.83 | 0.15 | 0.03 |
| BRN | Brunei Darussalam | Environment | 9 | 1.00 | 0.00 | 0.00 |
| BTN | Bhutan | Environment | 36 | 0.89 | 0.11 | 0.00 |
| BTN | Bhutan | HWB | 33 | 0.82 | 0.15 | 0.03 |
| BWA | Botswana | Environment | 33 | 0.88 | 0.03 | 0.09 |
| BWA | Botswana | HWB | 21 | 0.71 | 0.19 | 0.10 |
| CAF | Central African Republic | Environment | 66 | 0.92 | 0.08 | 0.00 |
| CAF | Central African Republic | HWB | 52 | 0.81 | 0.19 | 0.00 |
| CAN | Canada | Environment | 613 | 0.78 | 0.11 | 0.10 |
| CAN | Canada | HWB | 102 | 0.84 | 0.15 | 0.01 |
| CHE | Switzerland | Environment | 66 | 0.86 | 0.08 | 0.06 |
| CHE | Switzerland | HWB | 15 | 0.80 | 0.13 | 0.07 |
| CHL | Chile | Environment | 138 | 0.92 | 0.06 | 0.02 |
| CHL | Chile | HWB | 90 | 0.79 | 0.20 | 0.01 |
| CHN | China | Environment | 835 | 0.94 | 0.04 | 0.02 |
| CHN | China | HWB | 301 | 0.89 | 0.09 | 0.03 |
| CIV | Côte d’Ivoire | Environment | 27 | 0.89 | 0.07 | 0.04 |
| CIV | Côte d’Ivoire | HWB | 8 | 1.00 | 0.00 | 0.00 |
| CMR | Cameroon | Environment | 134 | 0.92 | 0.07 | 0.01 |
| CMR | Cameroon | HWB | 147 | 0.65 | 0.33 | 0.02 |
| COD | Democratic Republic of the Congo | Environment | 78 | 0.96 | 0.01 | 0.03 |
| COD | Democratic Republic of the Congo | HWB | 54 | 0.78 | 0.22 | 0.00 |

| | | | | | | |
|---|---|---|---|---|---|---|
| COG | Congo | Environment | 2 | 1.00 | 0.00 | 0.00 |
| COL | Colombia | Environment | 276 | 0.97 | 0.01 | 0.03 |
| COL | Colombia | HWB | 166 | 0.91 | 0.06 | 0.03 |
| CRI | Costa Rica | Environment | 242 | 0.93 | 0.02 | 0.05 |
| CRI | Costa Rica | HWB | 111 | 0.77 | 0.15 | 0.07 |
| CUB | Cuba | Environment | 26 | 0.85 | 0.08 | 0.08 |
| CUB | Cuba | HWB | 5 | 0.80 | 0.20 | 0.00 |
| CZE | Czech Republic | Environment | 26 | 0.69 | 0.23 | 0.08 |
| CZE | Czech Republic | HWB | 3 | 1.00 | 0.00 | 0.00 |
| DEU | Germany | Environment | 102 | 0.80 | 0.15 | 0.05 |
| DEU | Germany | HWB | 26 | 0.85 | 0.15 | 0.00 |
| DNK | Denmark | Environment | 10 | 0.90 | 0.00 | 0.10 |
| DNK | Denmark | HWB | 6 | 1.00 | 0.00 | 0.00 |
| DOM | Dominican Republic | Environment | 16 | 0.94 | 0.00 | 0.06 |
| DOM | Dominican Republic | HWB | 10 | 0.60 | 0.40 | 0.00 |
| DZA | Algeria | Environment | 21 | 0.95 | 0.00 | 0.05 |
| DZA | Algeria | HWB | 6 | 0.67 | 0.17 | 0.17 |
| ECU | Ecuador | Environment | 189 | 0.95 | 0.04 | 0.01 |
| ECU | Ecuador | HWB | 104 | 0.76 | 0.20 | 0.04 |
| EGY | Egypt | Environment | 3 | 1.00 | 0.00 | 0.00 |
| EGY | Egypt | HWB | 6 | 1.00 | 0.00 | 0.00 |
| ERI | Eritrea | Environment | 2 | 1.00 | 0.00 | 0.00 |
| ESP | Spain | Environment | 150 | 0.92 | 0.05 | 0.03 |
| ESP | Spain | HWB | 52 | 0.87 | 0.13 | 0.00 |
| EST | Estonia | Environment | 44 | 0.86 | 0.09 | 0.05 |
| EST | Estonia | HWB | 4 | 1.00 | 0.00 | 0.00 |
| ETH | Ethiopia | Environment | 87 | 0.94 | 0.02 | 0.03 |
| ETH | Ethiopia | HWB | 149 | 0.85 | 0.12 | 0.03 |
| FIN | Finland | Environment | 151 | 0.91 | 0.08 | 0.01 |
| FIN | Finland | HWB | 35 | 0.89 | 0.09 | 0.03 |
| FJI | Fiji | Environment | 8 | 1.00 | 0.00 | 0.00 |
| FJI | Fiji | HWB | 7 | 0.86 | 0.14 | 0.00 |
| FRA | France | Environment | 133 | 0.96 | 0.02 | 0.02 |
| FRA | France | HWB | 60 | 0.87 | 0.10 | 0.03 |
| GAB | Gabon | Environment | 27 | 0.89 | 0.11 | 0.00 |
| GAB | Gabon | HWB | 18 | 0.83 | 0.17 | 0.00 |
| GBR | U.K. of Great Britain and Northern Ireland | Environment | 133 | 0.83 | 0.14 | 0.04 |
| GBR | U.K. of Great Britain and Northern Ireland | HWB | 45 | 0.89 | 0.11 | 0.00 |
| GEO | Georgia | Environment | 30 | 0.90 | 0.07 | 0.03 |
| GEO | Georgia | HWB | 15 | 0.87 | 0.13 | 0.00 |
| GHA | Ghana | Environment | 98 | 0.94 | 0.04 | 0.02 |
| GHA | Ghana | HWB | 149 | 0.71 | 0.28 | 0.01 |
| GIN | Guinea | Environment | 14 | 0.71 | 0.29 | 0.00 |
| GIN | Guinea | HWB | 5 | 0.60 | 0.40 | 0.00 |
| GLP | Guadeloupe | HWB | 2 | 1.00 | 0.00 | 0.00 |
| GMB | Gambia | Environment | 3 | 1.00 | 0.00 | 0.00 |
| GMB | Gambia | HWB | 2 | 1.00 | 0.00 | 0.00 |
| GNB | Guinea-Bissau | Environment | 4 | 0.50 | 0.50 | 0.00 |
| GNQ | Equatorial Guinea | Environment | 17 | 1.00 | 0.00 | 0.00 |
| GNQ | Equatorial Guinea | HWB | 5 | 0.80 | 0.20 | 0.00 |
| GRC | Greece | Environment | 29 | 0.93 | 0.00 | 0.07 |
| GRC | Greece | HWB | 22 | 0.77 | 0.14 | 0.09 |
| GTM | Guatemala | Environment | 115 | 0.92 | 0.04 | 0.03 |
| GTM | Guatemala | HWB | 82 | 0.85 | 0.13 | 0.01 |

| | | | | | | |
|---|---|---|---|---|---|---|
| GUF | French Guiana | Environment | 2 | 1.00 | 0.00 | 0.00 |
| GUF | French Guiana | HWB | 2 | 0.50 | 0.00 | 0.50 |
| GUY | Guyana | Environment | 23 | 1.00 | 0.00 | 0.00 |
| GUY | Guyana | HWB | 15 | 0.80 | 0.20 | 0.00 |
| HND | Honduras | Environment | 61 | 0.98 | 0.00 | 0.02 |
| HND | Honduras | HWB | 54 | 0.76 | 0.24 | 0.00 |
| HRV | Croatia | Environment | 23 | 0.91 | 0.09 | 0.00 |
| HRV | Croatia | HWB | 1 | 1.00 | 0.00 | 0.00 |
| HTI | Haiti | Environment | 8 | 0.88 | 0.00 | 0.12 |
| HTI | Haiti | HWB | 8 | 0.88 | 0.12 | 0.00 |
| HUN | Hungary | Environment | 28 | 0.79 | 0.18 | 0.04 |
| HUN | Hungary | HWB | 8 | 1.00 | 0.00 | 0.00 |
| IDN | Indonesia | Environment | 821 | 0.95 | 0.03 | 0.02 |
| IDN | Indonesia | HWB | 680 | 0.79 | 0.19 | 0.02 |
| IND | India | Environment | 792 | 0.93 | 0.04 | 0.02 |
| IND | India | HWB | 891 | 0.81 | 0.18 | 0.01 |
| IRL | Ireland | Environment | 95 | 0.80 | 0.12 | 0.08 |
| IRL | Ireland | HWB | 6 | 0.83 | 0.17 | 0.00 |
| IRN | Iran (Islamic Republic of) | Environment | 145 | 0.89 | 0.05 | 0.06 |
| IRN | Iran (Islamic Republic of) | HWB | 59 | 0.92 | 0.08 | 0.00 |
| IRQ | Iraq | Environment | 7 | 1.00 | 0.00 | 0.00 |
| ISL | Iceland | Environment | 1 | 1.00 | 0.00 | 0.00 |
| ISL | Iceland | HWB | 4 | 0.75 | 0.25 | 0.00 |
| ITA | Italy | Environment | 215 | 0.85 | 0.13 | 0.01 |
| ITA | Italy | HWB | 61 | 0.87 | 0.13 | 0.00 |
| JOR | Jordan | Environment | 5 | 1.00 | 0.00 | 0.00 |
| JOR | Jordan | HWB | 6 | 1.00 | 0.00 | 0.00 |
| JPN | Japan | Environment | 130 | 0.85 | 0.12 | 0.04 |
| JPN | Japan | HWB | 83 | 0.83 | 0.13 | 0.04 |
| KAZ | Kazakhstan | Environment | 4 | 1.00 | 0.00 | 0.00 |
| KAZ | Kazakhstan | HWB | 4 | 0.25 | 0.75 | 0.00 |
| KEN | Kenya | Environment | 184 | 0.95 | 0.02 | 0.03 |
| KEN | Kenya | HWB | 134 | 0.83 | 0.16 | 0.01 |
| KGZ | Kyrgyzstan | Environment | 7 | 0.71 | 0.29 | 0.00 |
| KGZ | Kyrgyzstan | HWB | 4 | 1.00 | 0.00 | 0.00 |
| KHM | Cambodia | Environment | 75 | 0.97 | 0.00 | 0.03 |
| KHM | Cambodia | HWB | 87 | 0.83 | 0.11 | 0.06 |
| KOR | Republic of Korea | Environment | 49 | 0.76 | 0.18 | 0.06 |
| KOR | Republic of Korea | HWB | 32 | 0.97 | 0.03 | 0.00 |
| LAO | Lao People's Democratic Republic | Environment | 56 | 0.96 | 0.02 | 0.02 |
| LAO | Lao People's Democratic Republic | HWB | 94 | 0.73 | 0.22 | 0.04 |
| LBN | Lebanon | Environment | 1 | 0.00 | 0.00 | 1.00 |
| LBN | Lebanon | HWB | 2 | 1.00 | 0.00 | 0.00 |
| LBR | Liberia | Environment | 18 | 0.94 | 0.06 | 0.00 |
| LBR | Liberia | HWB | 13 | 1.00 | 0.00 | 0.00 |
| LKA | Sri Lanka | Environment | 82 | 0.89 | 0.07 | 0.04 |
| LKA | Sri Lanka | HWB | 56 | 0.71 | 0.29 | 0.00 |
| LTU | Lithuania | Environment | 51 | 0.78 | 0.18 | 0.04 |
| LTU | Lithuania | HWB | 1 | 1.00 | 0.00 | 0.00 |
| LUX | Luxembourg | Environment | 4 | 0.50 | 0.50 | 0.00 |
| LVA | Latvia | Environment | 6 | 0.67 | 0.33 | 0.00 |
| LVA | Latvia | HWB | 2 | 0.00 | 1.00 | 0.00 |
| MAR | Morocco | Environment | 25 | 0.96 | 0.04 | 0.00 |
| MAR | Morocco | HWB | 11 | 1.00 | 0.00 | 0.00 |

| | | | | | | |
|---|---|---|---|---|---|---|
| MDG | Madagascar | Environment | 124 | 0.90 | 0.04 | 0.06 |
| MDG | Madagascar | HWB | 129 | 0.50 | 0.48 | 0.02 |
| MEX | Mexico | Environment | 775 | 0.94 | 0.04 | 0.03 |
| MEX | Mexico | HWB | 421 | 0.81 | 0.16 | 0.03 |
| MLI | Mali | Environment | 3 | 0.67 | 0.33 | 0.00 |
| MLI | Mali | HWB | 4 | 0.00 | 1.00 | 0.00 |
| MLT | Malta | Environment | 2 | 1.00 | 0.00 | 0.00 |
| MMR | Myanmar | Environment | 81 | 0.96 | 0.01 | 0.02 |
| MMR | Myanmar | HWB | 76 | 0.74 | 0.18 | 0.08 |
| MNG | Mongolia | Environment | 1 | 1.00 | 0.00 | 0.00 |
| MNG | Mongolia | HWB | 5 | 0.80 | 0.20 | 0.00 |
| MOZ | Mozambique | Environment | 19 | 0.95 | 0.05 | 0.00 |
| MOZ | Mozambique | HWB | 17 | 0.76 | 0.24 | 0.00 |
| MRT | Mauritania | Environment | 2 | 1.00 | 0.00 | 0.00 |
| MTQ | Martinique | Environment | 1 | 1.00 | 0.00 | 0.00 |
| MWI | Malawi | Environment | 30 | 0.93 | 0.07 | 0.00 |
| MWI | Malawi | HWB | 31 | 0.90 | 0.10 | 0.00 |
| MYS | Malaysia | Environment | 236 | 0.94 | 0.03 | 0.03 |
| MYS | Malaysia | HWB | 106 | 0.81 | 0.17 | 0.02 |
| NAM | Namibia | Environment | 27 | 0.93 | 0.04 | 0.04 |
| NAM | Namibia | HWB | 22 | 0.91 | 0.09 | 0.00 |
| NCL | New Caledonia | Environment | 2 | 1.00 | 0.00 | 0.00 |
| NER | Niger | Environment | 7 | 1.00 | 0.00 | 0.00 |
| NER | Niger | HWB | 1 | 1.00 | 0.00 | 0.00 |
| NGA | Nigeria | Environment | 63 | 0.94 | 0.06 | 0.00 |
| NGA | Nigeria | HWB | 55 | 0.75 | 0.24 | 0.02 |
| NIC | Nicaragua | Environment | 45 | 0.96 | 0.02 | 0.02 |
| NIC | Nicaragua | HWB | 38 | 0.79 | 0.13 | 0.08 |
| NLD | Netherlands | Environment | 18 | 1.00 | 0.00 | 0.00 |
| NLD | Netherlands | HWB | 8 | 0.75 | 0.25 | 0.00 |
| NOR | Norway | Environment | 169 | 0.86 | 0.09 | 0.05 |
| NOR | Norway | HWB | 34 | 0.74 | 0.26 | 0.00 |
| NPL | Nepal | Environment | 340 | 0.92 | 0.06 | 0.01 |
| NPL | Nepal | HWB | 495 | 0.77 | 0.20 | 0.03 |
| NZL | New Zealand | Environment | 57 | 0.88 | 0.09 | 0.04 |
| NZL | New Zealand | HWB | 25 | 0.68 | 0.28 | 0.04 |
| PAK | Pakistan | Environment | 60 | 0.95 | 0.03 | 0.02 |
| PAK | Pakistan | HWB | 99 | 0.85 | 0.14 | 0.01 |
| PAN | Panama | Environment | 83 | 0.93 | 0.01 | 0.06 |
| PAN | Panama | HWB | 63 | 0.83 | 0.16 | 0.02 |
| PER | Peru | Environment | 162 | 0.97 | 0.02 | 0.01 |
| PER | Peru | HWB | 104 | 0.78 | 0.22 | 0.00 |
| PHL | Philippines | Environment | 88 | 0.92 | 0.07 | 0.01 |
| PHL | Philippines | HWB | 35 | 0.83 | 0.17 | 0.00 |
| PNG | Papua New Guinea | Environment | 43 | 0.95 | 0.05 | 0.00 |
| PNG | Papua New Guinea | HWB | 35 | 0.63 | 0.37 | 0.00 |
| POL | Poland | Environment | 97 | 0.75 | 0.16 | 0.08 |
| POL | Poland | HWB | 17 | 0.94 | 0.06 | 0.00 |
| PRK | Dem People's Rep of Korea | Environment | 2 | 1.00 | 0.00 | 0.00 |
| PRK | Dem People's Rep of Korea | HWB | 1 | 1.00 | 0.00 | 0.00 |
| PRT | Portugal | Environment | 44 | 0.89 | 0.02 | 0.09 |
| PRT | Portugal | HWB | 17 | 0.94 | 0.00 | 0.06 |
| PRY | Paraguay | Environment | 46 | 0.93 | 0.04 | 0.02 |
| PRY | Paraguay | HWB | 16 | 0.81 | 0.19 | 0.00 |

| | | | | | | |
|---|---|---|---|---|---|---|
| ROU | Romania | Environment | 25 | 0.72 | 0.12 | 0.16 |
| ROU | Romania | HWB | 11 | 1.00 | 0.00 | 0.00 |
| RUS | Russian Federation | Environment | 107 | 0.96 | 0.02 | 0.02 |
| RUS | Russian Federation | HWB | 19 | 0.84 | 0.16 | 0.00 |
| RWA | Rwanda | Environment | 17 | 1.00 | 0.00 | 0.00 |
| RWA | Rwanda | HWB | 14 | 1.00 | 0.00 | 0.00 |
| SDN | Sudan | Environment | 3 | 1.00 | 0.00 | 0.00 |
| SDN | Sudan | HWB | 2 | 1.00 | 0.00 | 0.00 |
| SEN | Senegal | Environment | 13 | 1.00 | 0.00 | 0.00 |
| SEN | Senegal | HWB | 12 | 1.00 | 0.00 | 0.00 |
| SLB | Solomon Islands | Environment | 4 | 1.00 | 0.00 | 0.00 |
| SLB | Solomon Islands | HWB | 3 | 1.00 | 0.00 | 0.00 |
| SLE | Sierra Leone | Environment | 4 | 1.00 | 0.00 | 0.00 |
| SLE | Sierra Leone | HWB | 1 | 1.00 | 0.00 | 0.00 |
| SLV | El Salvador | Environment | 20 | 1.00 | 0.00 | 0.00 |
| SLV | El Salvador | HWB | 13 | 0.92 | 0.08 | 0.00 |
| SOM | Somalia | Environment | 2 | 1.00 | 0.00 | 0.00 |
| SRB | Serbia | Environment | 6 | 0.83 | 0.17 | 0.00 |
| SRB | Serbia | HWB | 5 | 1.00 | 0.00 | 0.00 |
| SSD | South Sudan | Environment | 45 | 0.89 | 0.07 | 0.04 |
| SSD | South Sudan | HWB | 22 | 0.77 | 0.23 | 0.00 |
| STP | Sao Tome and Principe | Environment | 4 | 1.00 | 0.00 | 0.00 |
| SUR | Suriname | Environment | 13 | 1.00 | 0.00 | 0.00 |
| SUR | Suriname | HWB | 12 | 0.83 | 0.17 | 0.00 |
| SVK | Slovakia | Environment | 25 | 0.88 | 0.04 | 0.08 |
| SVK | Slovakia | HWB | 3 | 1.00 | 0.00 | 0.00 |
| SVN | Slovenia | Environment | 16 | 0.88 | 0.12 | 0.00 |
| SVN | Slovenia | HWB | 11 | 0.91 | 0.09 | 0.00 |
| SWE | Sweden | Environment | 95 | 0.86 | 0.05 | 0.08 |
| SWE | Sweden | HWB | 43 | 0.77 | 0.19 | 0.05 |
| SWZ | Swaziland | Environment | 4 | 0.75 | 0.25 | 0.00 |
| SWZ | Swaziland | HWB | 6 | 0.67 | 0.33 | 0.00 |
| SYR | Syrian Arab Republic | Environment | 15 | 0.93 | 0.07 | 0.00 |
| SYR | Syrian Arab Republic | HWB | 21 | 0.48 | 0.52 | 0.00 |
| TCD | Chad | HWB | 1 | 1.00 | 0.00 | 0.00 |
| TGO | Togo | Environment | 3 | 1.00 | 0.00 | 0.00 |
| TGO | Togo | HWB | 1 | 1.00 | 0.00 | 0.00 |
| THA | Thailand | Environment | 172 | 0.91 | 0.05 | 0.03 |
| THA | Thailand | HWB | 182 | 0.70 | 0.26 | 0.04 |
| TJK | Tajikistan | Environment | 1 | 1.00 | 0.00 | 0.00 |
| TJK | Tajikistan | HWB | 4 | 1.00 | 0.00 | 0.00 |
| TKM | Turkmenistan | Environment | 3 | 1.00 | 0.00 | 0.00 |
| TKM | Turkmenistan | HWB | 1 | 1.00 | 0.00 | 0.00 |
| TLS | Timor-Leste | Environment | 11 | 0.91 | 0.09 | 0.00 |
| TLS | Timor-Leste | HWB | 5 | 1.00 | 0.00 | 0.00 |
| TUN | Tunisia | Environment | 8 | 1.00 | 0.00 | 0.00 |
| TUN | Tunisia | HWB | 5 | 1.00 | 0.00 | 0.00 |
| TUR | Turkey | Environment | 51 | 0.88 | 0.08 | 0.04 |
| TUR | Turkey | HWB | 36 | 0.64 | 0.33 | 0.03 |
| TZA | United Republic of Tanzania | Environment | 201 | 0.91 | 0.08 | 0.01 |
| TZA | United Republic of Tanzania | HWB | 267 | 0.58 | 0.39 | 0.03 |
| UGA | Uganda | Environment | 162 | 0.90 | 0.04 | 0.06 |
| UGA | Uganda | HWB | 155 | 0.64 | 0.32 | 0.05 |
| UKR | Ukraine | Environment | 43 | 0.95 | 0.05 | 0.00 |

| | | | | | | |
|---|---|---|---|---|---|---|
| UKR | Ukraine | HWB | 7 | 0.86 | 0.14 | 0.00 |
| URY | Uruguay | Environment | 29 | 0.79 | 0.14 | 0.07 |
| URY | Uruguay | HWB | 13 | 0.62 | 0.31 | 0.08 |
| USA | United States of America | Environment | 2455 | 0.84 | 0.09 | 0.07 |
| USA | United States of America | HWB | 925 | 0.83 | 0.15 | 0.02 |
| UZB | Uzbekistan | Environment | 1 | 1.00 | 0.00 | 0.00 |
| VEN | Venezuela | Environment | 39 | 0.95 | 0.00 | 0.05 |
| VEN | Venezuela | HWB | 25 | 0.72 | 0.16 | 0.12 |
| VNM | Viet Nam | Environment | 121 | 0.91 | 0.04 | 0.05 |
| VNM | Viet Nam | HWB | 119 | 0.67 | 0.27 | 0.06 |
| YEM | Yemen | Environment | 8 | 1.00 | 0.00 | 0.00 |
| YEM | Yemen | HWB | 1 | 1.00 | 0.00 | 0.00 |
| ZAF | South Africa | Environment | 57 | 0.93 | 0.02 | 0.05 |
| ZAF | South Africa | HWB | 43 | 0.67 | 0.30 | 0.02 |
| ZMB | Zambia | Environment | 24 | 0.96 | 0.00 | 0.04 |
| ZMB | Zambia | HWB | 32 | 0.75 | 0.09 | 0.16 |
| ZWE | Zimbabwe | Environment | 68 | 1.00 | 0.00 | 0.00 |
| ZWE | Zimbabwe | HWB | 83 | 0.80 | 0.20 | 0.00 |

Table S5: The research attention and aboveground biomass of countries at highest risk of future forest conversion

| ISO3 | Name | Research attention | AGB | Quadrant |
|---|---|---|---|---|
| AGO | Angola | Data gap | Low | Emerging frontiers |
| ARG | Argentina | Mapped | High | Frontline hotspots |
| AUT | Austria | Data gap | High | Frontline hotspots |
| BDI | Burundi | Data gap | Med | Emerging frontiers |
| BEL | Belgium | Mapped | High | Frontline hotspots |
| BEN | Benin | Data gap | Med | Emerging frontiers |
| BFA | Burkina Faso | Data gap | Low | Emerging frontiers |
| BGD | Bangladesh | Data gap | Low | Frontline hotspots |
| BLR | Belarus | Data gap | High | Frontline hotspots |
| BLZ | Belize | Mapped | High | Frontline hotspots |
| BOL | Bolivia | Mapped | Med | Frontline hotspots |
| BRN | Brunei Darussalam | Mapped | High | Emerging frontiers |
| CIV | Côte d'Ivoire | Data gap | High | Frontline hotspots |
| CMR | Cameroon | Mapped | High | Frontline hotspots |
| COD | Democratic Republic of the Congo | Data gap | High | Frontline hotspots |
| CRI | Costa Rica | Mapped | High | Emerging frontiers |
| CUB | Cuba | Data gap | Med | Frontline hotspots |
| DNK | Denmark | Data gap | Med | Frontline hotspots |
| DOM | Dominican Republic | Data gap | Med | Frontline hotspots |
| DZA | Algeria | Data gap | Low | Frontline hotspots |
| ECU | Ecuador | Mapped | High | Emerging frontiers |
| ESP | Spain | Mapped | Low | Frontline hotspots |
| EST | Estonia | Mapped | Med | Frontline hotspots |
| FIN | Finland | Mapped | Low | Frontline hotspots |
| GHA | Ghana | Mapped | Med | Frontline hotspots |
| GIN | Guinea | Data gap | Med | Frontline hotspots |
| GLP | Guadeloupe | Data gap | High | Emerging frontiers |
| GMB | Gambia | Data gap | Low | Emerging frontiers |
| GNB | Guinea-Bissau | Data gap | Low | Frontline hotspots |
| GNQ | Equatorial Guinea | Mapped | High | Emerging frontiers |

| | | | | |
|---|---|---|---|---|
| GTM | Guatemala | Mapped | High | Frontline hotspots |
| HND | Honduras | Mapped | Low | Frontline hotspots |
| HTI | Haiti | Data gap | Med | Frontline hotspots |
| IDN | Indonesia | Mapped | High | Frontline hotspots |
| IRL | Ireland | Mapped | Med | Frontline hotspots |
| JOR | Jordan | Data gap | Low | Emerging frontiers |
| KEN | Kenya | Mapped | Med | Emerging frontiers |
| KHM | Cambodia | Mapped | Med | Frontline hotspots |
| LAO | Lao People's Democratic Republic | Mapped | Med | Frontline hotspots |
| LBR | Liberia | Mapped | High | Frontline hotspots |
| LKA | Sri Lanka | Mapped | Low | Emerging frontiers |
| LUX | Luxembourg | Mapped | High | Frontline hotspots |
| LVA | Latvia | Mapped | Med | Frontline hotspots |
| MDG | Madagascar | Mapped | Med | Frontline hotspots |
| MLI | Mali | Data gap | Med | Emerging frontiers |
| MMR | Myanmar | Data gap | Med | Frontline hotspots |
| MOZ | Mozambique | Data gap | Low | Frontline hotspots |
| MRT | Mauritania | Data gap | Low | Emerging frontiers |
| MTQ | Martinique | Data gap | High | Emerging frontiers |
| MWI | Malawi | Data gap | Low | Frontline hotspots |
| MYS | Malaysia | Mapped | High | Frontline hotspots |
| NGA | Nigeria | Data gap | Med | Frontline hotspots |
| NIC | Nicaragua | Mapped | Med | Frontline hotspots |
| NLD | Netherlands | Data gap | High | Emerging frontiers |
| NOR | Norway | Mapped | Low | Emerging frontiers |
| PAN | Panama | Mapped | High | Emerging frontiers |
| PHL | Philippines | Data gap | High | Emerging frontiers |
| POL | Poland | Data gap | High | Frontline hotspots |
| PRT | Portugal | Mapped | Low | Frontline hotspots |
| PRY | Paraguay | Mapped | Low | Frontline hotspots |
| RWA | Rwanda | Data gap | High | Frontline hotspots |
| SEN | Senegal | Data gap | Low | Frontline hotspots |
| SLE | Sierra Leone | Data gap | Med | Frontline hotspots |
| SLV | El Salvador | Mapped | Med | Emerging frontiers |
| SWE | Sweden | Mapped | Low | Frontline hotspots |
| SWZ | Swaziland | Mapped | Med | Frontline hotspots |
| TCD | Chad | Data gap | Low | Frontline hotspots |
| TGO | Togo | Data gap | Low | Frontline hotspots |
| THA | Thailand | Mapped | High | Frontline hotspots |
| TUN | Tunisia | Data gap | Low | Frontline hotspots |
| TZA | United Republic of Tanzania | Mapped | Low | Frontline hotspots |
| UGA | Uganda | Mapped | Med | Frontline hotspots |
| URY | Uruguay | Mapped | Med | Frontline hotspots |
| VNM | Viet Nam | Data gap | Low | Frontline hotspots |
| ZMB | Zambia | Mapped | Low | Frontline hotspots |
| ZWE | Zimbabwe | Mapped | High | Frontline hotspots |

### 1.4 Evaluating potential signatures of publication bias

We investigated the degree of evidence for publication bias potentially distorting our results. We did so by first evaluating the distribution of positive, neutral, and negative evidence as a function of two abstract attributes: the citation count and the journal's selectivity. Citation counts reflect the amount of amplification of a finding among the research community. Therefore, we sought to see if there were clear patterns where the least or most amplified abstracts tended to have systematically more or less positive, negative, or neutral evidence. Ultimately, for the 20,625 abstracts with citation count data, we did not observe any clear patterns for negative, neutral, or positive evidence increasing or decreasing across citation count quartiles (Figure S4). Instead, we saw broad concordance in the overall makeup of the evidence across citation count categories, with differences within 1-2%.

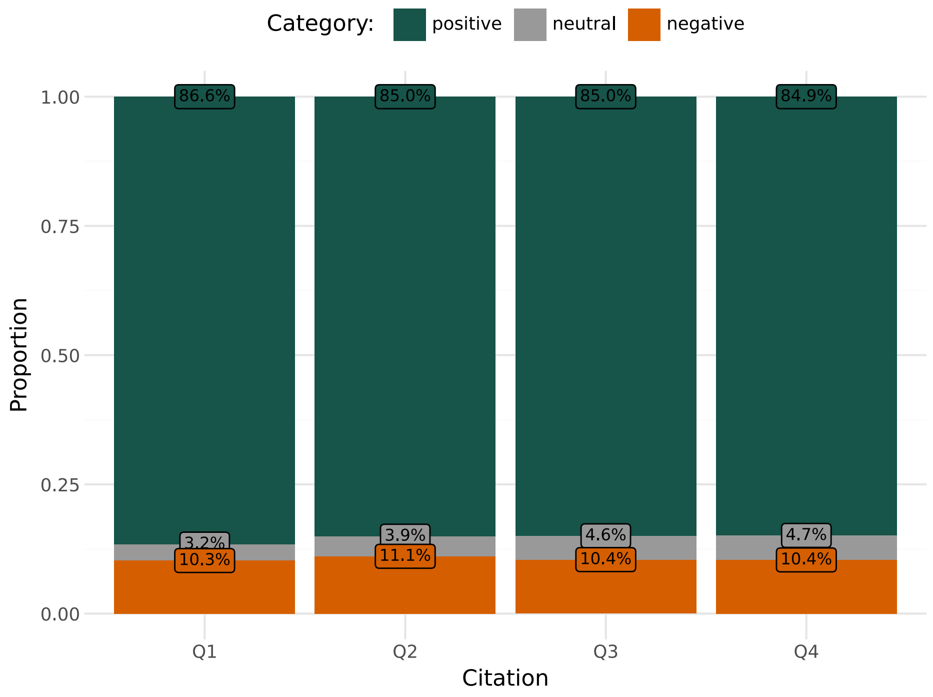


Figure S4: The distribution of positive, neutral, and negative evidence as a function of abstract citation count. The citation count increases along the x-axis from the 1st to the 4th quartile (Q1: (0, 4] citations, n=6,012 abstracts, Q2: (4, 11] citations, n=4,376 abstracts, Q3: (11, 28] citations, n=5,124 abstracts, Q4: (28, 1628] citations, n=5,113 abstracts).

Pivoting to journal selectivity, we joined our AFC evidence mapping dataset with SCImago Journal Rank (SJR) data based on ISSN. We observed non-null and non-missing SJR values for 8,663 abstracts. To proxy journal selectivity, we used the SJR Best Quartile metric, which assigns the top quartile (Q4-Q1 in descending order) to a journal based on its best performance (e.g., if a journal is ranked Q1 for discipline A but Q3 for discipline B, its SJR Best Quartile metric would be Q1). Initially, we saw that there was a highly imbalanced count of Q1-Q4 journals ($n_{q1} = 5790, n_{q2} = 1524, n_{q3} = 822, n_{q4} = 518$). Hence, we combined the least selective quadrats into a Q3+ category. As before, we did not see a pattern where the most or least selective journals exhibited systematically greater or lower positive evidence (Figure S5). While Q2 journals exhibited the most distinct pattern of positive, neutral, and negative evidence, we saw values within 1-3% for all directional categories between Q1 and Q3+ journals.

Subsequently, we sought to evaluate if differential patterns in publishing between high versus low-income countries and across different continents could be distorting our results. To conduct a rapid assessment, we performed a mixed-effects logistic regression (using `lme4` in `R`):

$$p_{pos} \sim \zeta_{\text{country}} + \beta_{\text{domain (HWB or Environment)}} + \beta_{\text{region}} + \beta_{\text{Income}} + \beta_{FPP} + \beta_{pop_{2019}}$$

In this model, $FPP$ corresponded to the proportion of a country's population that was considered forest-proximate (residing within 5km of a forest, (*30*)), while $pop_{2019}$ was the overall population size in 2019, scaled across countries.

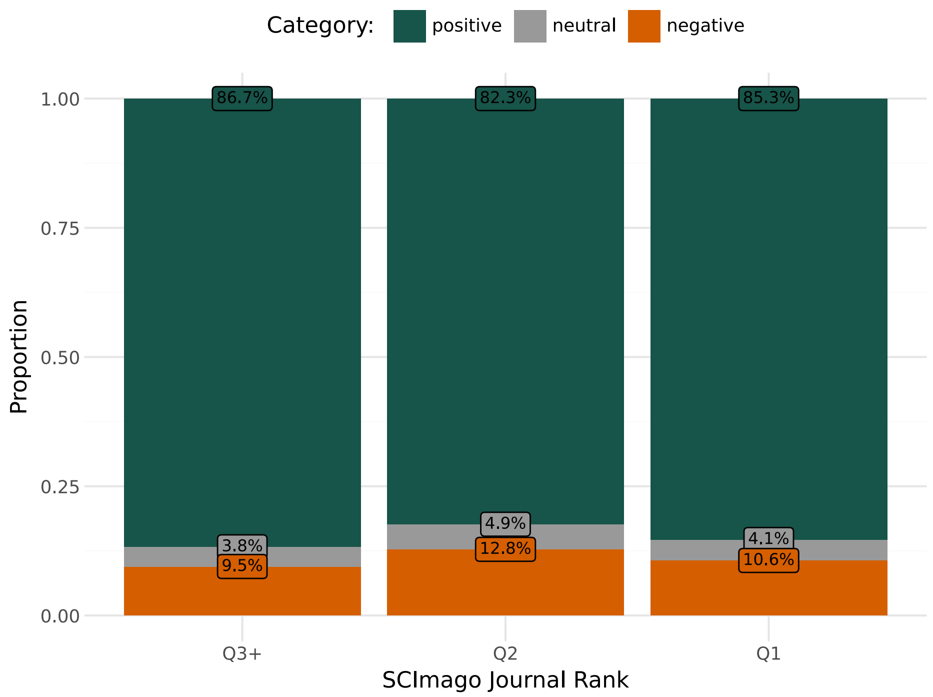


Figure S5: The distribution of directional AFC co-impact evidence as a function of journal selectivity, proxied by SJR Best Quartile rank. Typically, journal selectivity increases with impact factor, or rank. The x-axis displays increasing journal selectivity from Q3+ (n=1,340 abstracts) to Q2 (n=1,524 abstracts) to Q1 (n=5,790 abstracts).

Based on existing literature, one could expect that lower-income countries, and those with a larger $FPP$, might report more trade-offs for AFC and thus more negative evidence. Similarly, one might expect that continents with a larger number of lower-income and tropical countries may also report greater trade-offs with AFC. From past bibliometric research, one might also expect that such countries publish at lower rates, and that this could artificially skew our evidence base to be more positive than warranted. However, we observed that *higher income* countries in our dataset were associated with slightly more negative evidence for AFC, contravening our expectation; importantly, we did not observe that high income countries were underrepresented in our dataset. We did not observe any significant, clearly directional effects of any continent on the degree of positive AFC evidence, nor did we observe any clear association between $FPP$ and AFC evidence. As such, our data suggest that the degree of positive AFC co-benefit evidence is not an artifact of different publication rates as a function of geography or reduced country-level income.

Table S6: Robustness check using a mixed effects regression of positive evidence versus overall evidence across 136 countries with complete data.

| | Coefficient estimates | |
|---|---|---|
| Variable | Coefficient | 95% CI |
| Domain: HWB | $-1.325^{**}$ | ($-1.414$, $-1.237$) |
| Low Income | 0.262 | ($-0.247$, 0.772) |
| Low-Middle Income | 0.376 | ($-0.030$, 0.782) |
| Upper-Middle Income | $0.456^{*}$ | (0.104, 0.808) |
| Region: Asia | 0.012 | ($-0.302$, 0.325) |
| Region: Europe | 0.215 | ($-0.265$, 0.694) |
| Region: North and Central America | 0.245 | ($-0.162$, 0.653) |
| Region: Oceania | 0.018 | ($-0.583$, 0.619) |
| Region: South America | 0.270 | ($-0.153$, 0.693) |
| Forest-proximate population share | $-0.025$ | ($-0.922$, 0.872) |
| Overall population | 0.046 | ($-0.031$, 0.122) |
| Observations | 269 | |
| Log Likelihood | $-650.933$ | |
| Akaike Inf. Crit. | 1,327.867 | |
| Bayesian Inf. Crit. | 1,374.598 | |
| *Note:* | $^{*}$p<0.05; $^{**}$p<0.01 | |

31. Food and Agriculture Organization of the United Nations, "FAO. Global Forest Resources Assessment 2020", tech. rep. (UN FAO - processed by Our World in Data, 2026).
